# TMAO Destabilizes RNA Secondary Structure via Direct Hydrogen Bond Interactions

**DOI:** 10.1101/2022.08.03.502647

**Authors:** Samuel S. Cho, Adam T. Green, Changbong Hyeon, D. Thirumalai

## Abstract

Trimethylamine N-oxide (TMAO) is an osmolyte that accumulates in cells in response to osmotic stress. TMAO stabilizes proteins by the entropic stabilization mechanism, which pictures TMAO as a nano-crowder that predominantly destabilizes the unfolded state. However, the mechanism of action of TMAO on RNA is much less understood. Here, we use all atom molecular dynamics simulations to investigate how TMAO interacts with a 12-nt RNA hairpin with a high melting temperature, and an 8-nt RNA hairpin, which has a relatively fluid native basin in the absence of TMAO. The use of the two hairpins with different free energy of stabilization allows us to probe the origin of the destabilization effect of TMAO on RNA molecules without the possibility of forming tertiary interactions. We generated multiple trajectories using allatom molecular dynamics (MD) simulations in explicit water by employing AMBER and CHARMM force fields, both in the absence and presence of TMAO. We observed qualitatively similar RNA-TMAO interaction profiles from the simulations using the two force fields. TMAO hydrogen bond interactions are largely depleted around the paired RNA bases and ribose sugars. In contrast, we show that the oxygen atom in TMAO, the hydrogen bond acceptor, preferentially interacts with the hydrogen bond donors in the solvent exposed bases, such as those in the stem-loop, the destabilized base stacks in the unfolded state, especially in the marginally stable 8-nt RNA hairpin. The predicted destabilization mechanism through TMAO-RNA hydrogen bond interactions could be tested using two-dimensional IR spectroscopy. Since TMAO does not significantly interact with the hydroxyl group of the ribose sugars, we predict that similar results must also hold for DNA.

## Introduction

Osmolytes comprise a class of naturally occurring simple organic compounds used by cells to tune the relative stability of biomolecules such as proteins and RNA in response to osmotic stress.^1–4^ The increase in the concentration of some stabilizing osmolytes, such as Trimethylamine-N-oxide (TMAO), can increase the thermodynamic stability and enzymatic activity of biological molecules by biasing their structures forwards the folded state, thereby maintaining their viability in harsh environments without compromising their function(s). This is to be contrasted with the accumulation of destabilizing agents, such as urea or guanidinium chloride, which promote denaturation so that their function(s) becomes diminished or abolished.

The importance of the interactions between TMAO with proteins and RNA cannot be overstated. A few reasons are: (1) It has been discovered that bacteria in the gut breakdown L-carnitine (found in red meat for example) producing trimethyl amine, which is converted to TMAO in the liver. Elevated levels of TMAO are apparently correlated with high platelet hyper-reactivity and potential for heart diseases.^5,6^ (2) It has been shown that TMAO increases the *in vitro* assembly of the 50S ribosomal subunit from rRNA and proteins by hundred fold.^7^ (3) Experiments show that TMAO destabilizes (marginally) secondary structures of RNA^4^ while stabilizing tertiary interactions.^3,4^ The explanation of the enhanced stability of tertiary structures of RNA is nuanced, and could depend on the presence of a fraction of TMAO molecules that are protonated,^8^ thus giving it a positive charge.

The underlying molecular mechanism by which osmolytes stabilize or destabilize a biomolecule has been the subject of investigation for a number of years.^9–11^ In contrast to RNA, TMAO-induced enhancement in the stability of proteins is reasonably understood. Our proposal that TMAO entropically stabilizes the folded state by restricting the conformational fluctuations of the unfolded state by behaving as a nano-crowder mechanism is supported by experiments by Gai and coworkers.^12,13^ The nano-crowder effect could originate by plausible formation of cluster-like structure between TMAO and a few water molecules, as proposed using experiments^14^ and computer simulations.^15^

The “indirect mechanism” suggests that TMAO alters and rearranges the surrounding water molecules,^14^ and the ordering of a network of the water molecules stabilizes the folded state of the protein by decreasing the entropy of the unfolded state. ^15^ It follows then that the destabilization of the water structure by a destabilizing osmolyte would impede hydrophobic collapse and destabilize the folded state. ^16^ However, recent studies point to a “direct mechanism” for osmolyte-induced stabilization and destabilization of biomolecules. ^11,17^ In this view, the osmolyte interacts directly with the biomolecules to affect their stabilities. For proteins, these interactions are widely thought to involve specific interactions with predominantly the protein backbone ^9,17^ and to a lesser extent the side chains. ^17^ The stability of the native state results from the depletion of osmolytes^18^ from the surface of proteins. If the osmolytes interact directly with the surface of the biomolecule, the folded state would be destabilized at sufficiently high osmolyte concentrations. In an earlier study, Cho et al. proposed an entropic stabilization mechanism^17^ whereby depleted TMAO “crowd” the protein and limit the fluctuations of the unfolded state, thereby stabilizing the folded state. ^19^ Experiments by Gai and coworkers have validated our prediction that TMAO acts as a nano-crowder using 2D IR spectroscopy. ^12^

Although less studied, the effect of TMAO on the ground state of RNA has been investigated experimentally and theoretically.^4,8,20,21^ In a key paper by Lambert and Draper, the effect of different osmolytes for a number of RNA molecules was carried out. In all cases, TMAO stabilized the tertiary interactions. Interestingly, a small destabilization of the secondary structure was observed. ^4^ To complement these studies, Denning et al. predicted, using atomistic MD simulations, that TMAO induced stabilization of RNA tertiary structure could be pH dependent. They showed that for PreQ1 riboswitch, TMAO destabilizes tertiary structure but its protonated form stabilizes it. ^8^ In atom MD simulations combined with coarse-grained models of guanosine-5’-monophosphate in the presence of TMAO, Pincus et al. identified significant hydrogen bond interactions with the N2 atom of an amide moiety that could destabilize the secondary strutures of RNA. ^20^

In the present study, we performed AMBER and CHARMM all-atom MD simulations of a stable 12-nt and an unstable 8-nt RNA hairpin, which are ideal model systems of folded and unfolded secondary structures. We systematically identified specific RNA-TMAO interactions by quantifying the interactions using radial distribution functions of the hydrogen bond interactions in the 12-nt and 8-nt hairpins. In accord with experiments, we found that TMAO destabilizes the hairpins because of the enhanced hydrogen bond interaction between the hydrogen atom on the carbonyl of TMAO with certain amide nitrogen atoms on the RNA bases. It is possible to test our predictions using two-dimensional IR spectroscopy experiments.

## Results

### Unstable and Stable RNA Hairpins Represent Unfolded and Folded States

To understand the RNA-TMAO interactions in the folded and unfolded states, we performed AMBER and CHARMM all-atom MD simulations of two RNA hairpins: a 12-nt (Fig. 1a,b) and an 8-nt (Fig. 1c, d) RNA hairpin. These two RNA hairpins were selected on the basis of their stabilities. The 12-nt hairpin is extremely stable with an unusually high melting temperature.^22^ In contrast, for the 8-nt RNA hairpin the nature of the native ensemble is under investigation because of the unusual kinetics observed in the folding of the hairpin. Gruebele and coworkers showed that the folding rate increased with temperature,^23^ which is inconsistent with the folding kinetics of a protein with a unique, well-defined native state. Replica-exchange all-atom simulations by Garcia and coworkers suggested that the native ensemble is composed of multiple hairpin structures.^24^ Throughout our simulations, the 12-nt hairpin maintains the native structure with the base-pairs largely intact while the 8-nt hairpin partially unfolds and possibly reforms native base-pair interactions (Fig. S1,S2).

**Figure 1:**
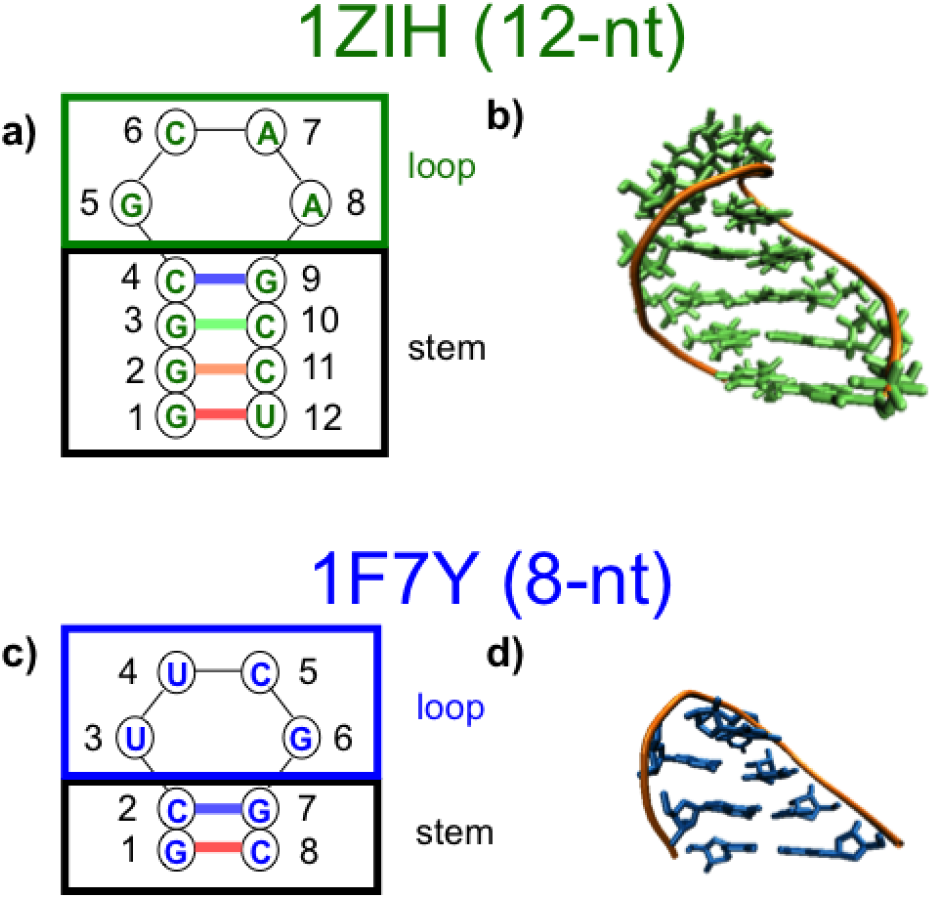
Schematic of the secondary (left) and the ground state hairpin (right) structures of the 12-nt (a,b) and 8-nt (c,d) hairpins simulated in our study. In the secondary structures, each colored thick lines denotes a base-pair.

We performed simulations using both AMBER and CHARMM force fields. The TMAO parameters developed by Kast et al. were largely taken from the CHARMM force field as a starting point, but the electrostatics parameters were calculated using AMBER.^25^ It is well known that nucleic acids generally have stability issues in both the AMBER and CHARMM force fields, which could lead to unrealistic trapped states.^26^ For example, Chen and García tuned the van der Waals parameters of the AMBER force field to reproduce the folding of three RNA tetraloops within 1-3 Å RMSD,^27^ possibly allowing the accurate folding of larger RNA. Here, we compare the simulation results of two widely used and independently developed force fields and assess the qualitative similarities and differences of the simulated RNA secondary structures and their interactions with TMAO. Because of potential inaccuracies in the force fields our results are likely to be only qualitative.

### TMAO Destabilizes Secondary Structure of a Stable Hairpin

For the stable 12-nt RNA hairpin, we performed two sets of three independent 1 *μs* simulations, one in the absence of TMAO and another in the presence of 1.0 M TMAO with the AMBER99-bs0 and CHARMM36 force fields. To quantify the extent of secondary structure formed in the hairpin, we calculated the distances between base pairs. In the absence of TMAO, the simulations of the hairpin (Fig. S1) yielded a very stable configuration with the average base-to-base distances of about 6 Å that largely did not fluctuate throughout the entire MD simulations with the AMBER99-bs0 force field. For the CHARMM36 force field MD simulations, only the terminal GU base pair, which is non-Watson-Crick (non-WC), exhibited transient increases that is significantly beyond 6 Å in the two trajectories (Fig. S1e,f), and the hairpin began to unfold after about 700 ns in one trajectory (Fig. S1d).

We repeated the MD simulations except in 1.0 M TMAO solution, and observed qualitatively similar results (Fig. S2) in the AMBER99-bs0 force field MD simulations (Fig. S2a-c). Results from two trajectories were nearly identical to those observed in the absence of TMAO with all base-to-base distances at around 6 Å (Fig. S1a-c). However, the distance between base pair 4-9 (near the loop) increased significantly to more than 10 Å. In addition, the distance between base pair (1-12) increases to more than 20 Å in one trajectory after about 550 ns (Fig. S2b), suggesting that the hairpin reorganized into a different structure.

On the other hand, the base-to-base distances (average distance is about 6 Å) throughout the three CHARMM 36 force field MD simulation trajectories (Fig. S2d-f) show that the base pairs were largely maintained. However, there are substantial fluctuations in the base-to-base distances, ranging between about 2-9 Å for the 2-11, 3-10, and 4-9 base pairs and more than 20 Å for the 1-12 base pair in simulations without TMAO (Fig. S1d-f). In both of these sets of trajectories with TMAO, the RNA hairpin becomes destabilized by adopting a different structure (Fig. S2b) or having greater fluctuations of their base-pairs.

### TMAO is largely depleted around paired RNA bases and ribose sugars

To quantify the TMAO interactions with the RNA hairpin, we calculated the radial distribution functions (RDFs) between the oxygen atom in TMAO (the only possible hydrogen bond acceptor), and the various hydrogen bond donors in the RNA hairpins in the 1.0 M-TMAO. We first compared the interactions between TMAO and the RNA bases and sugars (Fig. 2). For both AMBER99-bs0 and CHARMM36, we observe only modest peaks at about 3 Å (Fig. 2a,b), which would correspond to the optimal hydrogen bonding distance. The RDF differences of the 12-nt hairpin loop and stem for TMAO-Base interactions were minimal except that the RDF peaks of AMBER99-bs0 MD simulations are slightly greater than those of CHARMM36 ones (Fig. 2a), and they were negligible for TMAO-Sugar interactions. While all ribose sugar interactions are chemically identical in RNA, they were distinct in bases.

**Figure 2:**
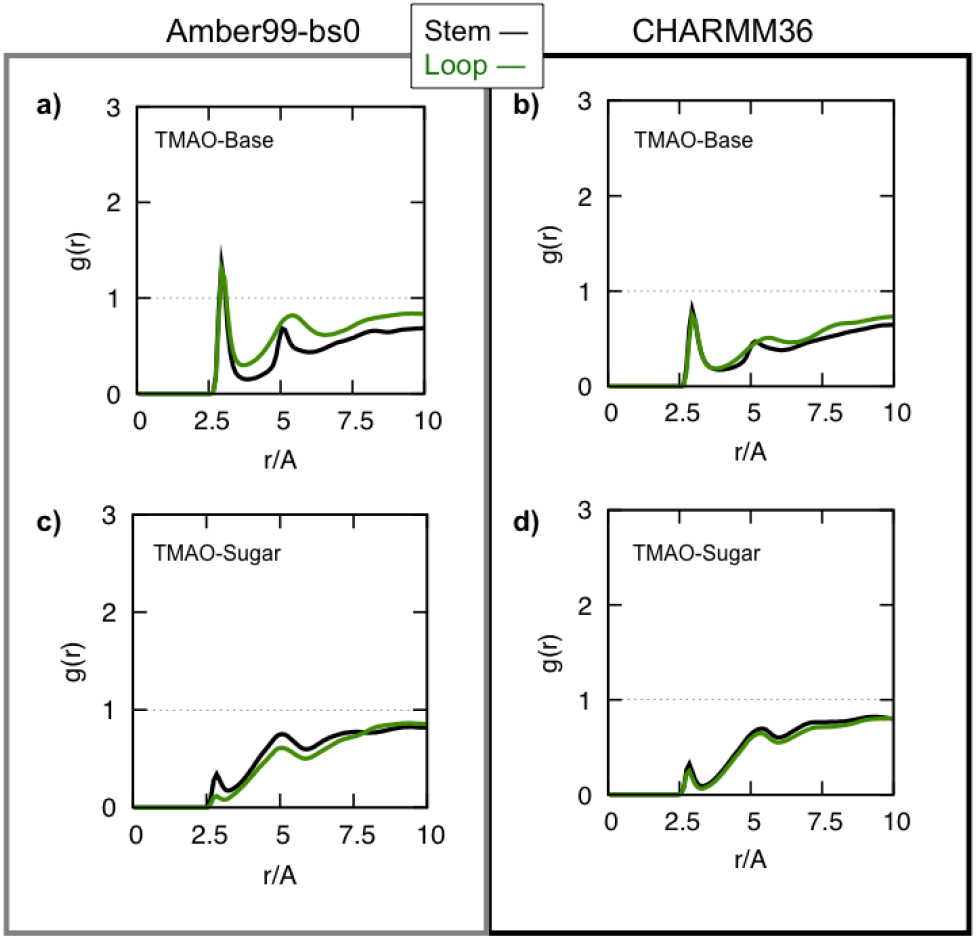
Radial distribution function between the TMAO oxygen and all the RNA base possible hydrogen bond donors (a,b) and ribose sugar possible hydrogen bond donors (c,d) for the loop nucleotides and stem nucleotides in the 12-nt RNA hairpin.

We then calculated the RDFs for the specific interactions of TMAO with every possible individual hydrogen bond donors in bases (Fig. 3). A significant peak in the RDF is seen for interactions of TMAO with the stem guanine amide moiety (N2) (Fig. 3d-f) but not amine (N1) (Fig. 3a-c) and loop cytosine amide moiety (N4) (Fig. 3g-i). Interestingly, the adenine in the loop also has an amide moiety, but its peak was only modestly significant even though it is expected to be solvent exposed (Fig. 3j-l). In the stable RNA hairpin, the stem hydrogen bond donors, with the exception of the terminal GU pair, are occupied with base-pair interactions, and do not directly interact with TMAO via hydrogen bonding. Lambert et al. ^4^ suggested phosphate dehydration in the presence of TMAO to explain their experimental observations, which we do not observe in either the AMBER or the CHARMM force field MD simulations (Fig. S3).

**Figure 3:**
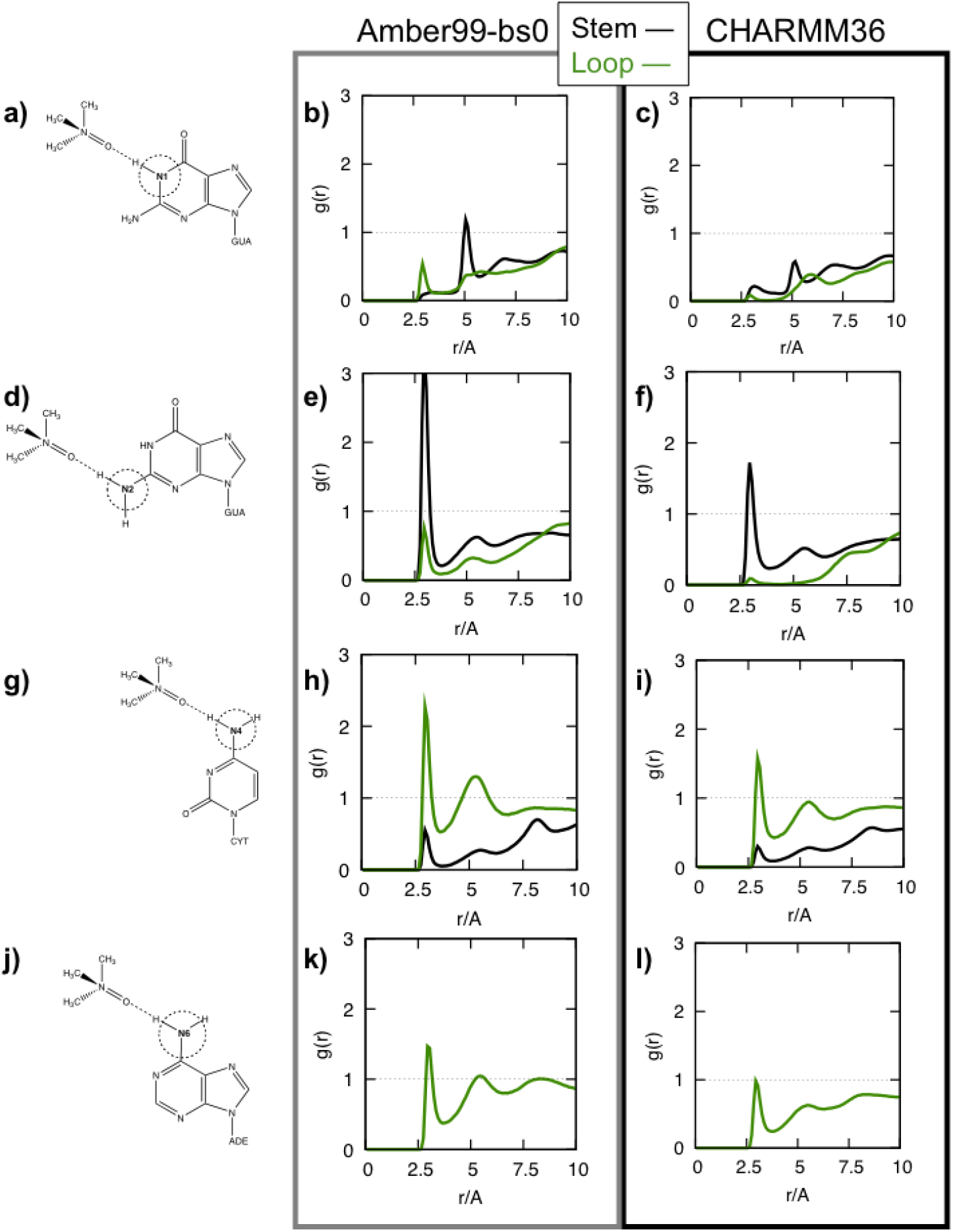
Radial distribution function between the TMAO oxygen and the individual possible specific RNA base hydrogen bond donors: (a-c) guanine N1, (d-f) guanine N2, (g-i) cytosine N4, and (j-l) adenine N6 in the 12-nt RNA hairpin. The leftmost column shows the TMAO interaction studied, and the center and right columns are the radial distribution functions plotted for the AMBER (center) and CHARMM (right) MD simulations.

### TMAO Interacts Directly with the Bases of the Unstable RNA Hairpin via Hydrogen Bonds

We then performed MD simulations of the unstable 8-nt RNA hairpin and observed a more complicated behavior in its dynamics. In the absence of TMAO, the AMBER99-bs0 force field MD simulation of the 8-nt hairpin was very stable in one trajectory (Fig. S5b), similar to the analogous MD simulations of the 12-nt hairpin (Fig. S1a-c), but in two other trajectories exhibited unfolding and refolding (Fig. S5a,c). However, for the CHARMM36 force field MD simulations, the base-to-base distances deviated significantly from around 6 Å and dissolved the base pairs such that the hairpins were essentially unfolded throughout the trajectories (Fig. S5d-f). We repeated the MD simulations except with 1.0 M TMAO, and we observed qualitatively similar results (Fig. S5).

The RDFs between TMAO and the possible hydrogen bond donors in the bases yielded a modest peak at about 3 Å for the stems in the bases (Fig. 4a,b), but only a negligible peak was observed for the sugar hydrogen bond donors (Fig. 4c,d). Furthermore, the most pronounced peaks in the RDFs were the stem guanine amide moiety (N2) but not amine (N1) for both AMBER99-bs0 and CHARMM36 (Fig. 5a-f) and only a modest peak in the loop cytosine amide moiety (N4) (Fig. 5g-i). A modest peak is observed for the loop uracil amide (N3) that is slightly more pronounced in CHARMM36.

**Figure 4:**
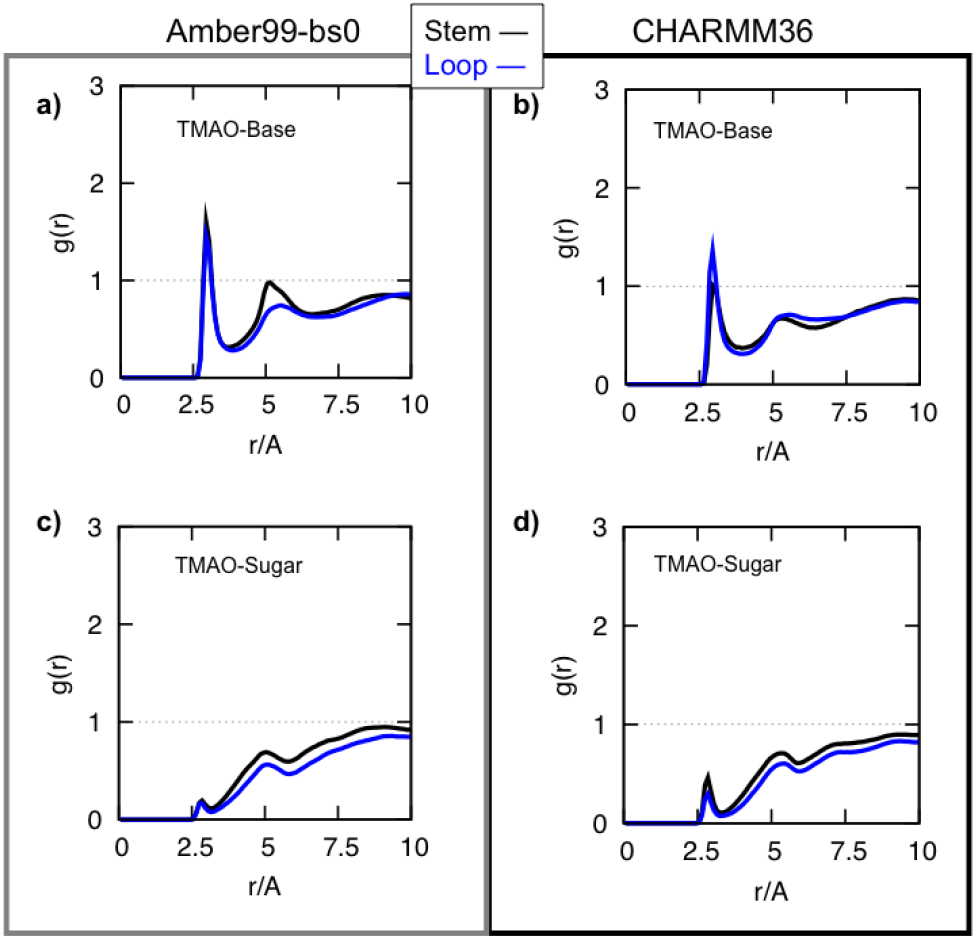
Radial distribution function between the TMAO oxygen and all the RNA base possible hydrogen bond donors (a,b) and ribose sugar possible hydrogen bond donors (c,d) for the loop nucleotides and stem nucleotides in the 8-nt RNA hairpin.

**Figure 5:**
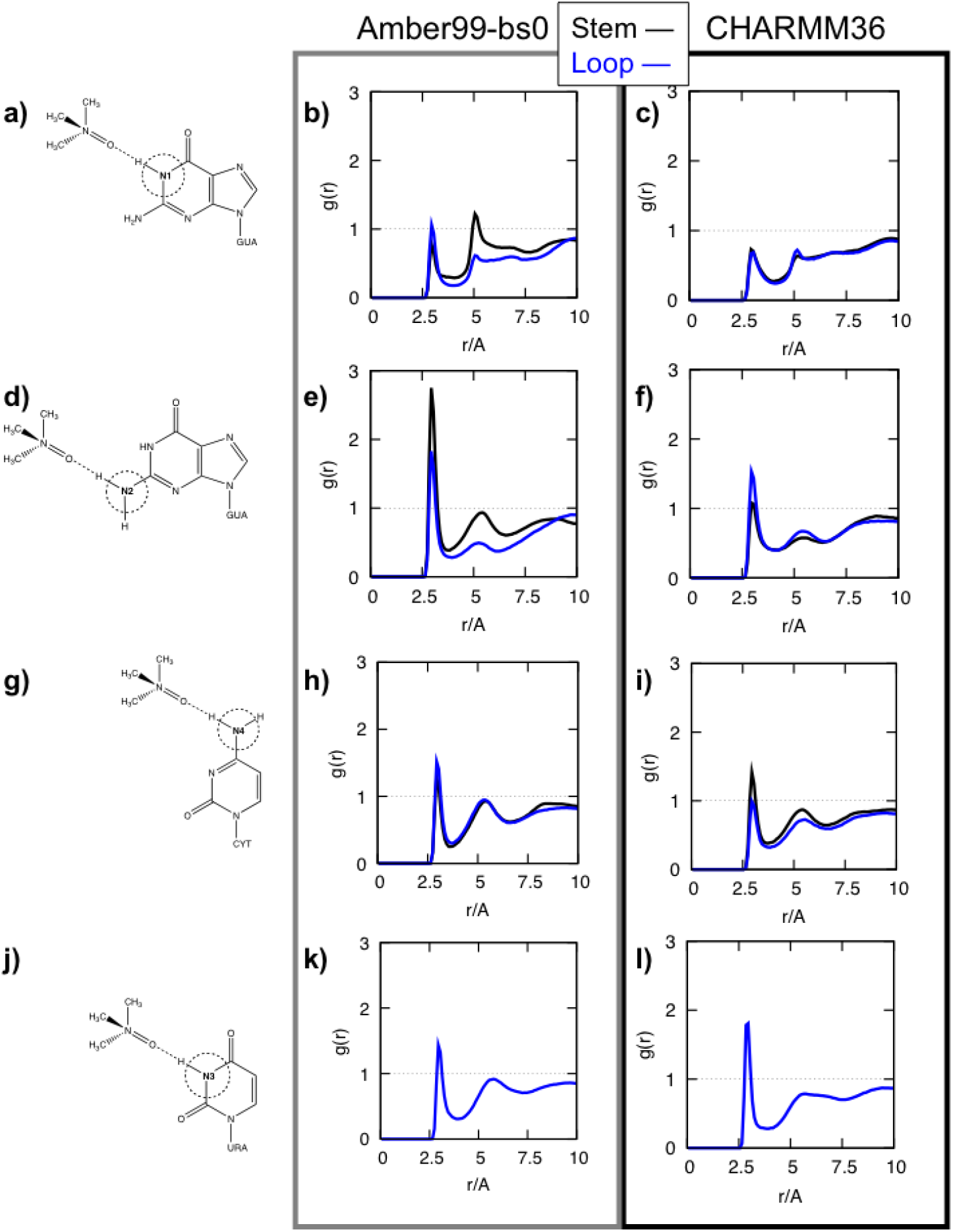
Radial distribution function between the TMAO oxygen and the individual possible specific RNA base hydrogen bond donors: (a-c) guanine N1, (d-f) guanine N2, (g-i) cytosine N4, and (j-l) uracil N3 in the 8-nt RNA hairpin. The leftmost column shows the TMAO interaction studied, and the center and right columns are the radial distribution functions plotted for the AMBER (center) and CHARMM (right) MD simulations.

## Discussion

To identify the specific interactions between TMAO and RNA and characterize the impact on RNA secondary structure, we performed all atom molecular dynamics simulations using AMBER and CHARMM empirical force fields using two model RNA hairpins that represent the folded and unfolded states, respectively: a stable 12-nt hairpin and an unstable 8-nt hairpin. As reported in previous studies, the secondary structure is more stable in AMBER than in CHARMM. In both the force fields, we observe that TMAO is largely depleted around the stable folded 12-nt hairpin whereas TMAO interacts directly with the bases of the unstable 8-nt hairpin with solvent exposed bases. This is consistent with an entropic stabilization mechanism for osmolyte-induced folding that we had previously for proteins. Furthermore, in both the force fields we observe qualitatively consistent hydrogen bond interactions between TMAO and the RNA hairpins. TMAO interacts through preferential hydrogen bonds with the amide moieties of the solvent exposed bases. In agreement with the previous CHARMM MD simulation study of Pincus et al.,^20^ the N2 in guanine, in particular, had the most significant interactions with TMAO as quantified by the radial distribution functions. There are also other hydrogen bond interactions with solvent exposed nucleotide bases. Cumulatively, our study shows that direct hydrogen bond interactions between the carbonyl oxygen of TMAO with the bases destabilizes the secondary structures modestly disrupting the hydrogen bond formation between base pairs.

### Contrasting the destabilization mechanism of Urea and TMAO

Previously we showed that urea destabilizes the 22 nucleotide RNA hairpin (P5GA) by engaging in hydrogen bonds with the bases.^28^ The nitrogen and oxygen atoms of urea are engaged in multiple hydrogen bond interactions with the bases as well as the phosphate groups. Detailed examination of the structures established that urea forms stacking type interactions, which strongly disrupts the Watson-Crick bases pairs. A very similar mechanism was discovered in the course of accounting for disruption of tertiary structures in PreQ1 riboswitch at high urea concentrations.^29^

In contrast, only the carbonyl oxygen of TMAO is involved in direct hydrogen bond interactions with nitrogen atoms in the bases. The inability to form multiple hydrogen bonds with RNA readily explains the much weaker effect of TMAO on the stability of RNA hairpins. Our finding, that TMAO is depleted near the bases and sugar moieties also suggests that tertiary interactions interactions are likely stabilized by nano-crowding effects, as was proposed for proteins,^17^ and subsequently validated using 2D infrared spectroscopy.^12^

### TMAO avoids proteins but interacts favorably with RNA hairpins

In a previous study probing the action of TMAO on polypeptide chains, ^17^ it was shown that there are hydrogen bond interactions between the osmolyte and the peptides. However, the predominant stabilizing effect of TMAO on peptide of any length is the depletion of the solute from the protein surface. This allows for maximization of intra protein interaction. The most vivid illustration of this effect is the formation of helix formation in Aβ_16–22_, which is random coil in isolation.^30,31^ Furthermore, Muttathukattil and Reddy recently showed that TMAO aids in fibril growth of the Sup35 prion peptide by stabilizing the locked state.^32^ In sharp contrast, TMAO forms hydrogen bond interactions with RNA bases, which disrupts modestly the hairpin structures. Thus, TMAO *avoids* the surface of proteins whereas it forms *favorable* interactions with RNA.

## Conclusion

The observation that marine organisms have evolved to use TMAO to counteract osmotic stress in the oceans is used to rationalize the importance of studying the effects of osmolytes on proteins. Needless to say that RNA also must function under those conditions. Nevertheless, the physical basis of the stabilization or destabilization mechanism on RNA has not been carefully investigated. We have shown that direct hydrogen bond formation between TMAO and RNA results in mild destabilization of RNA hairpins. Because the predicted effect is small (see Lambert and Draper^4^), we suggest that 2D IR would be an ideal method, as was done in model peptides. ^12^

## Methods

### TMAO Force Fields

To investigate how RNA hairpins are affected by the presence of TMAO, we performed MD simulations using the NAMD program suite^33^ and the AMBER99-bs0^34^ and CHARMM36^35^ force fields the for nucleic acids, waters, and ions. The TMAO force field parameters of Kast et al. ^25^ were used for simulations with the AMBER and CHARMM force fields. There exist several other variants of TMAO force field parameters that are meant to reproduce some experimental observations. ^36^ Rodriguez-Ropero et al. evaluated four different TMAO force field parameters on model of a hydrophobic synthetic homopolymer, including the one used here, and observed that the general features of TMAO for all force fields were largely similar at concentrations till about 1M TMAO. ^37^ Because we are only interested in elucidating the general mechanism of TMAO action on RNA secondary structures, we believe that the small differences in the force field are not relevant in elucidating the destabilization mechanism. In any event, there are no studies of TMAO effects using the different force fields examined in the context of polymers and small proteins.^36,37^

### Choice of RNA Hairpins

We chose two RNA hairpins: a stable 12-nt RNA hairpin with a high melting temperature, and a relatively unstable 8-nt RNA hairpin. The crystal structure of the isolated 12-nt hairpin is known (PDB ID: 1ZIH).^22^ The X-ray crystallographic structure of the isolated 8-nt hairpin has not been resolved, although its kinetics has been studied experimentally, ^23^ so we truncated the 8-nt hairpin from the crystal structure of an RNA tetraloop (PDB ID: 1F7Y)^38^ for our simulations.

### MD Simulations

We performed two sets of all-atom MD simulations with periodic boundary conditions, one in the presence of TMAO and another in the absence of TMAO, for each RNA hairpin. As a starting point, the RNA molecules were centered in a rectangular water box comprised of TIP3P water molecules, and all water molecules within 1.6 Å of the RNA molecules were deleted. The dimension of the water box was set to be equal to 10 Å more than the longest length of the RNA molecule. We also added sodium ions to neutralize the phosphate ions on the RNA backbone. The ions positions were set using the MEADIONIZE plugin module. ^39^ The TMAO molecules were randomly placed into the system by replacing water molecules until the desired 1.0 M concentration was obtained. For each RNA simulation, the initial configuration was energy minimized for 2,000 steps using the NAMD conjugate gradient module, and only the lowest energy configuration was as the starting coordinates for simulations. The system was then equilibrated by performing 50 ps MD simulations while increasing the temperature to its target 298 K. Analyses were performed for the production run trajectories after equilibration.

We then performed three independent simulations of each RNA hairpin at 298 K in the absence and presence of TMAO, using both the AMBER and CHARMM force fields, resulting in twelve (2 x 3 simulations for each force field) 1 *μs* trajectories for each hairpin. The distances between the center of mass distances of the base pairs were calculated over each set of trajectories. The radial distribution function was calculated over each set of three independent simulations between the TMAO oxygen and each of the hydrogen bonding donors in the RNA hairpins except when indicated.

## Acknowledgement

This work was supported in part by a grant from the National Science Foundation (CHE19-00033) and the Welch Foundation through the Collie-Welch chair (F-0019). We also acknowledge computational support from the Wake Forest University DEAC Cluster, where the MD simulations were performed.

## Supporting Information Available

A listing of the contents of each file supplied as Supporting Information should be included. For instructions on what should be included in the Supporting Information as well as how to prepare this material for publications, refer to the journal’s Instructions for Authors.

The following files are available free of charge.

- Filename: TMAO-RNA-supplementary-fig.pdf

## Notes

### Competing Interest Statement

The authors have declared no competing interest.

