## Supplementary Information for "TMAO Destabilizes RNA Secondary Structure via Direct Hydrogen Bond Interactions"

### 1ZIH (12-nt) w/o TMAO

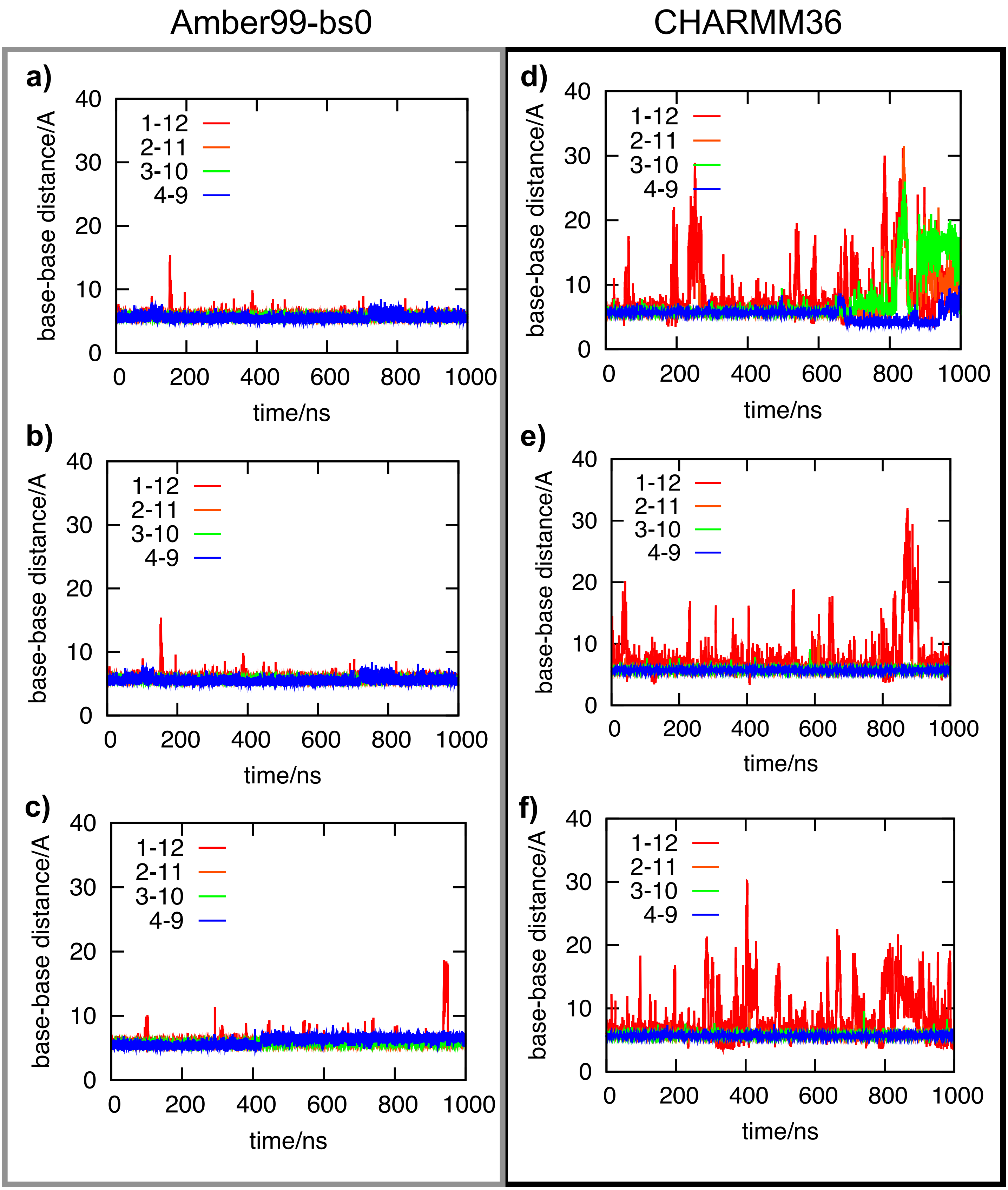


Fig. S1: Cho et al.

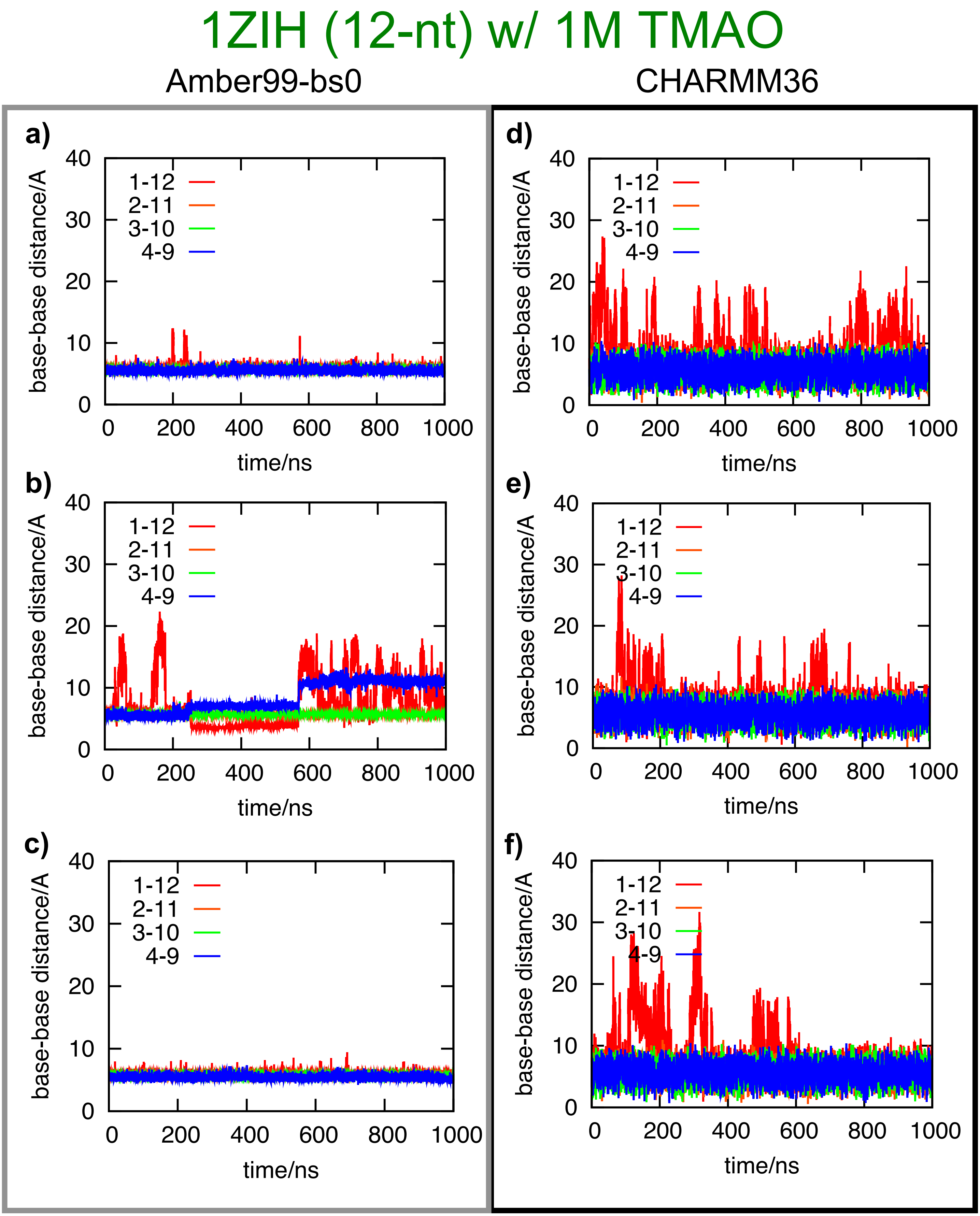


Fig. S2: Cho et al.

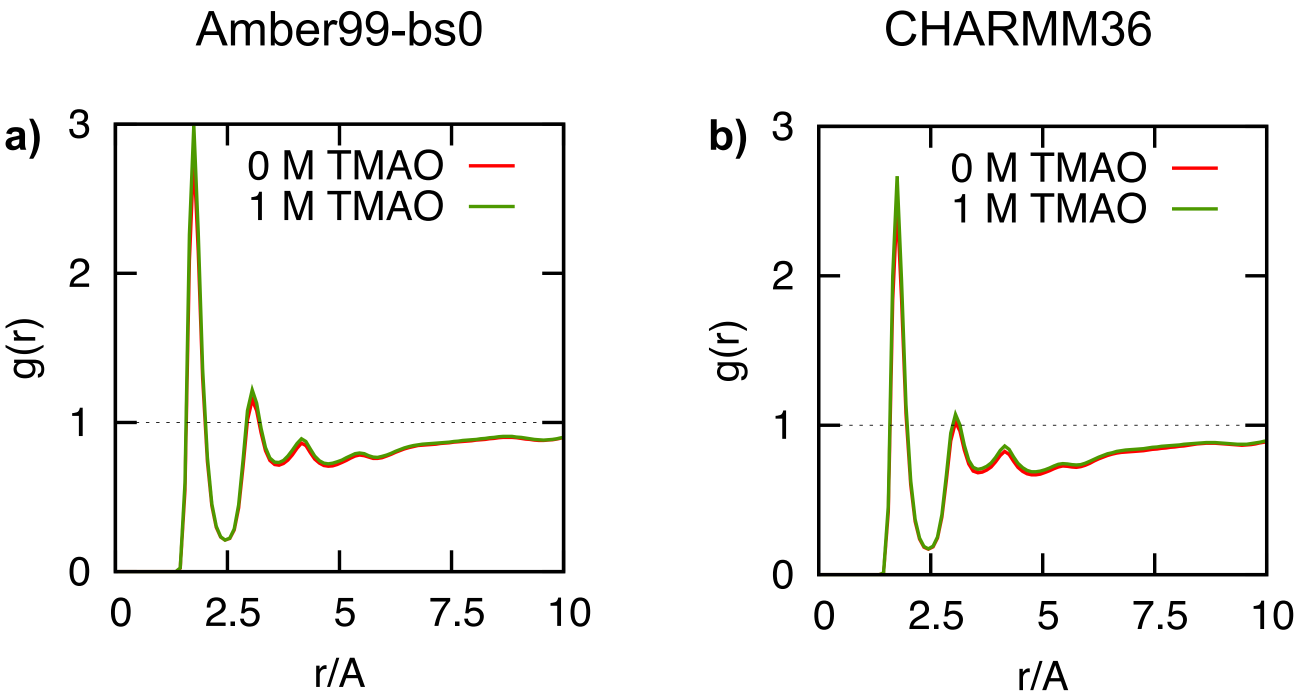


Fig. S3: Cho et al.

### 1F7Y (8-nt) w/o TMAO

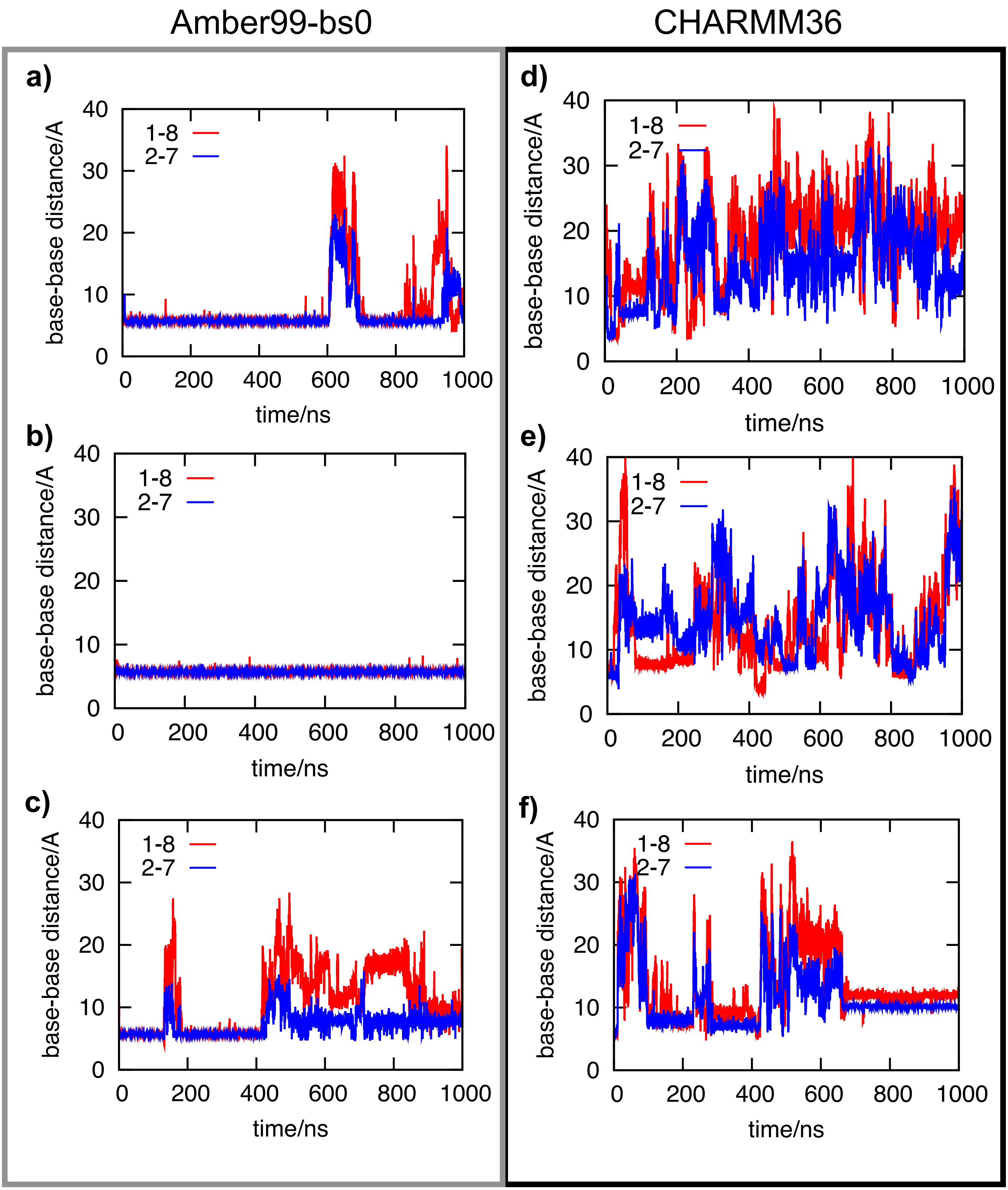


Fig. S4: Cho et al.

### 1F7Y (8-nt) w/ 1M TMAO

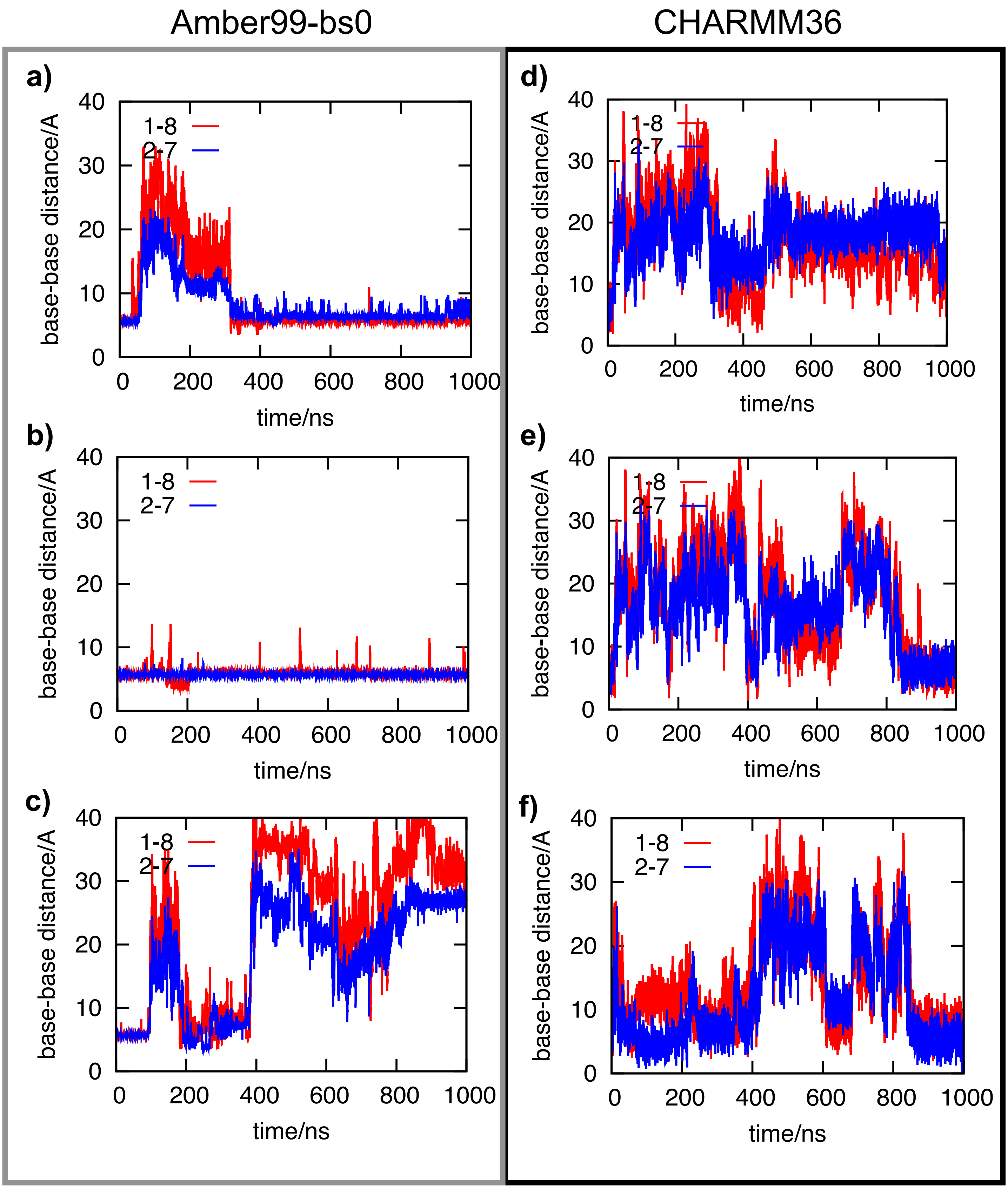


Fig. S5: Cho et al.
